# Frontal plane ankle stiffness increases with axial load independent of muscle activity

**DOI:** 10.1101/2021.12.06.471410

**Authors:** Zoe Villamar, Eric J. Perreault, Daniel Ludvig

## Abstract

Ankle sprains are the most common musculoskeletal injury, typically resulting from excessive inversion of the ankle. One way to prevent excessive inversion and maintain ankle stability is to generate a stiffness that is sufficient to resist externally imposed rotations. Frontal-plane ankle stiffness increases as participants place more weight on their ankle, but whether this effect is due to muscle activation or axial loading of the ankle is unknown. Identifying whether and to what extent axial loading affects ankle stiffness is important in understanding what role the passive mechanics of the ankle joint play in maintaining its stability. The objective of this study was to determine the effect of passive axial load on frontal-plane ankle stiffness. We had subjects seated in a chair as an axial load was applied to the ankle ranging from 10% to 50% body weight. Small rotational perturbations were applied to the ankle in the frontal plane to estimate stiffness. We found a significant, linear, 3-fold increase in ankle stiffness with axial load from the range of 0% bodyweight to 50% bodyweight. This increase could not be due to muscle activity as we observed no significant axial-load-dependent change in any of the recorded muscle activations. These results demonstrate that axial loading is a significant contributor to maintaining frontal-plane ankle stability, and that disruptions to the mechanism mediating this sensitivity of stiffness to axial loading may result in pathological cases of ankle instability.

## INTRODUCTION

Ankle sprains are the most common musculoskeletal injury, accounting for at least 3 million hospital visits per year in the United States (Doherty et al., 2014). Nearly 85% of sprains occur due to excess movement of the ankle in the frontal plane, specifically in inversion (Andersen et al., 2004). The ankle requires sufficient stability in the frontal plane to avoid excessive inversion and minimize the likelihood of sprains. The contribution of unloaded passive structures and muscle activation to maintaining ankle stability in the frontal plane has been well documented (Ashton-Miller et al., 1996; Leardini et al., 2000). However, cadaveric studies suggest that axial-loading-sensitive mechanisms are a major contributor to ankle stability during weight-bearing conditions. Any contribution to stability by axial loading could be very important as ankle experiences up to 2–3 times body weight during many locomotor activities (Firminger et al., 2018). Identifying the role of axial loading in maintaining ankle stability is important to determine whether an impairment in this mechanism could be a causative factor in recurring ankle sprains.

While frontal-plane ankle stability has been investigated during weight-bearing conditions, none have isolated the contributions to stability made by axial loading of the ankle. Ankle stability has been quantified by computing the impedance, the dynamic relationship between an imposed movement and the resistive torque generated in response (Kearney and Hunter, 1990). The impedance has often been modeled as a spring-mass-damper system quantified by its stiffness, inertia, and viscosity respectively (Ludvig and Kearney, 2007; Rouse et al., 2014), with the stiffness component being the widely studied component. During standing, axial load due to weight-bearing has been shown to increase frontal plane stiffness, but this increase in axial load was accompanied by an increase in muscle activation (Matos et al., 2021). Muscle activation is known to substantially increase ankle stiffness (Lee et al., 2014a; Lee et al., 2014b; Mirbagheri et al., 2000). Thus, it remains unknown what the effect of axial loading independent of muscle activity is to human ankle stability.

Cadaveric studies have suggested that axial load may affect frontal-plane ankle stability, but the extent to which these studies extend to *in vivo* ankle stability is unclear. Under axial load transmitted via the tibia, the ankle undergoes less rotation in inversion/eversion, rotation in the horizontal plane, and flexion for a given torque (McCullough and Burge, 1980; Stiehl et al., 1993; Stormont et al., 1985). This reduction in range of motion for a fixed torque is equivalent to an increase in ankle stiffness (McCullough and Burge, 1980; Stiehl et al., 1993). However, these cadaveric studies are limited. There are two joints within the ankle joint complex that allow motion in the frontal plane: the talocrural joint and the subtalar joint (Hertel, 2002). These cadaveric studies isolated motion to the talocrural joint (Stiehl et al., 1993; Watanabe et al., 2012), discounting rotation at the subtalar joint, where most of the inversion occurs under *in vivo* conditions (Arndt et al., 2004). Additionally, these studies remove much of the surrounding soft tissues, including muscle and tendon, thus altering the biomechanics of the ankle. While cadaver studies are insightful in that they provide evidence that load may affect ankle stability, they do not conclusively describe how an *in vivo* ankle behaves under axial load.

The objective of this study was to determine the effect of passive axial load on frontal-plane ankle stiffness. We hypothesized that ankle stiffness would increase with an increase in passive axial load. We had subjects seated in a chair as an axial load was applied to the ankle ranging from 10% to 50% body weight through the controlled application of force on the knee. Small rotational perturbations were applied to the ankle in the frontal plane to estimate stiffness. We tested our hypothesis by determining if there was a significant increase in ankle stiffness with an increase in axial load. To ensure that any changes in stiffness could only be attributed to the passive axial load, we tested whether there was a relationship between the axial load and the muscle activity of any of the surrounding muscles. If ankle stiffness is sensitive to passive axial loading, it would suggest that the mechanism responsible for this sensitivity to passive loading may substantially contribute to ankle stability during functional weight-bearing conditions.

## METHODS

### Participants

Fifteen individuals (8 males, 7 females; age = 27.2 ± 4.4 (mean ± standard deviation), body mass: 75 ± 14 kg, height: 1.74 ± 0.10 m) participated in this study. All participants had no known neurological or connective tissue disorder. Participants were excluded if they had a known orthopeadic diagnosis, bone fracture in the lower extremity that required realignment, or history of significant ankle sprains in the past 5 years, with significant being defined as resulting in a loss of at least one day of desired physical activity. Each participant was right foot dominant and each participant’s right ankle was tested. Ethical approval for the study was received from the Northwestern University Institutional Review Board (STU00009204 and STU00215245). Written informed consent was obtained prior to testing.

### Experimental Set-up

To quantify changes in frontal-plane ankle stiffness due to imposed axial loading, we applied an axial load on their leg and perturbed their ankle as they were seated. Each participant was seated in an adjustable chair (Biodex Medical Systems, Inc. Shirley, NY, USA). To apply perturbations directly to the ankle, we rigidly secured the participant’s foot to an electric rotary motor (BSM90N-3150AF, Baldor, Fort Smith, AR, USA) via a custom-made fiberglass cast (**Figure 1**). Knee angle and ankle angle were both set at 90° of flexion using a goniometer. Straps were tightened around the torso, waist, and right leg to limit movements during the experiment. To apply the axial load to the ankle, a C-shaped pad positioned just proximal to the knee joint was lowered by the experimenter. The desired load was set by the experimenter, aided by visual feedback of the axial force from the load cell located underneath the participant’s foot. Ankle angle was simultaneously recorded using a 24-bit quadrature encoder card (PCI-QUAD04, Measurement Computing, Norton, MA, USA). A six-degree-of-freedom load cell (45E15A4, JR3, Woodland, CA, USA) measured all ankle forces and torques. Data acquisition and control of the rotary motor were executed using xPC target (The Mathworks Inc., Natick, MA).

**Figure 1.**
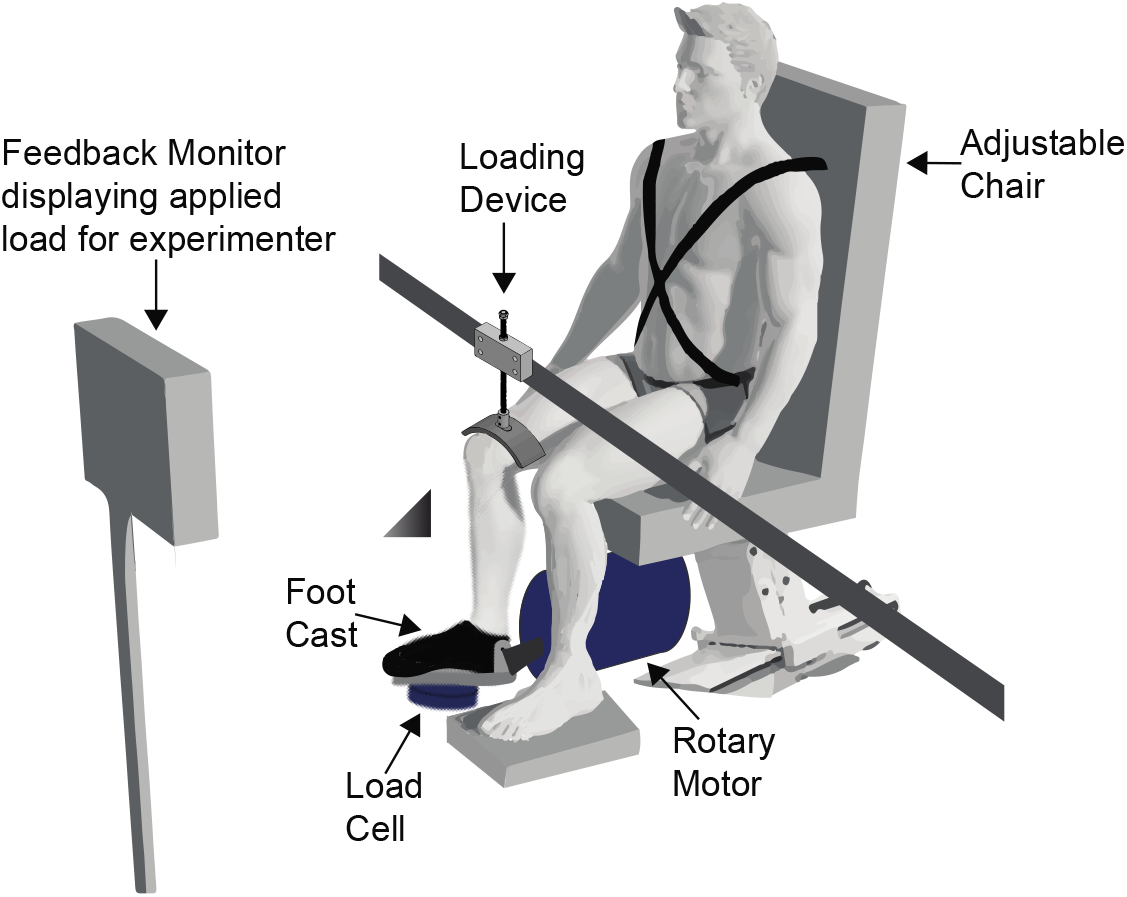
Experimental Set-Up. Participants were seated in an adjustable chair and had their ankles fixed to a load cell and rotary motor via a custom-made foot cast. External axial loading was applied via a loading device placed above the knee. This load was adjusted by the experimenter, aided visually by a feedback monitor displaying the applied axial load, as measured by the load cell.

Electromyography (EMG) data were collected to ensure the muscle activity did not change with axial loading. EMGs were collected using surface electrodes (Bagnoli, Delsys Inc, Boston, MA) from the following muscles: tibialis anterior, lateral gastrocnemius, medial gastrocnemius, soleus, peroneus longus, and peroneus brevis. These muscles were chosen because they surround the ankle and can contribute to the torque generated in the frontal and sagittal planes (Brockett and Chapman, 2016). Standard skin preparation methods were used before adhering each electrode to the skin (Besomi et al., 2019; Merletti and Muceli, 2019). EMG measurements were amplified (Delsys Bagnoli, Natick, MA) as needed to maximize the range of the data acquisition system. All analog data were passed through an anti-aliasing filter (500 Hz using a 5-pole Bessel filter) and sampled at 2.5 kHz (PCI-DAS1602/16, Measurement Computing, Norton, MA, USA).

### Protocol

Initial maximum voluntary contraction (MVC) trials were first collected to normalize the EMG signals. Participants were asked to complete maximum voluntary contractions in 6 directions: inversion/eversion, dorsiflexion/plantarflexion, and internal/external rotation. Each maximum voluntary contraction was repeated twice. EMG from these trials were later used to normalize the EMG to allow for comparison across participants. A resting trial in which the participant was instructed to relax without the application of load or perturbations was also collected.

The study was designed to determine the effect of passive load on ankle stiffness. Participants were instructed to remain relaxed during each trial. Axial loads were applied up to a maximum of 50% bodyweight (BW). We chose maximum of 50% BW since that was near the comfort limit of pressure for the participants. The unloaded condition refers to the participant being seated in the setup without the external load applied; thereby only the weight of the foot and shank were recorded in this condition. Across all participants, this load was 10.9 ± 1.3% body weight (mean ± standard deviation). The loading conditions were unloaded, 20% BW, 30% BW, 40% BW, 50% BW. Each loading condition was repeated twice. The order of the trials was randomized. To estimate the frontal-plane ankle stiffness, we applied rotational perturbations in the frontal plane. The perturbations were a pseudorandom binary sequence of amplitude 0.03 radians, 3.0 rad/s, and 0.150 switching time (Matos et al., 2021). Each trial lasted 65 seconds. We also estimated the impedance of the equipment and cast with a trial in which only the cast was perturbed. All measures of impedance were normalized by the weight of the participant.

### Quantifying Ankle Impedance

We quantified the stability of the ankle in the frontal plane by computing its impedance. First, we decimated the data to 50 Hz. Then, we characterized the impedance of the ankle by computing a non-parametric impulse response function (IRF) between the measured ankle angle and the measured torque in the frontal plane (**Figure 2**) (Matos et al., 2021). To compute the stiffness, viscosity, and inertia from the impedance, we first inverted the impedance to the admittance, and then fit a parametric 2^nd^ order model to the admittance IRF (Kearney et al., 1997; Ludvig and Perreault, 2011). Separately, we computed the stiffness, viscosity, and inertia of the cast and apparatus from the cast-only trial, and removed these measures from the respective measures of the experimental trials. For the experimental trials, the parametric 2^nd^ order model accounted for 92.2 ± 7.7% (mean ± standard deviation) of the measured torque. We performed all data processing and analysis in MATLAB (Mathworks, Natick, MA, USA).

**Figure 2.**
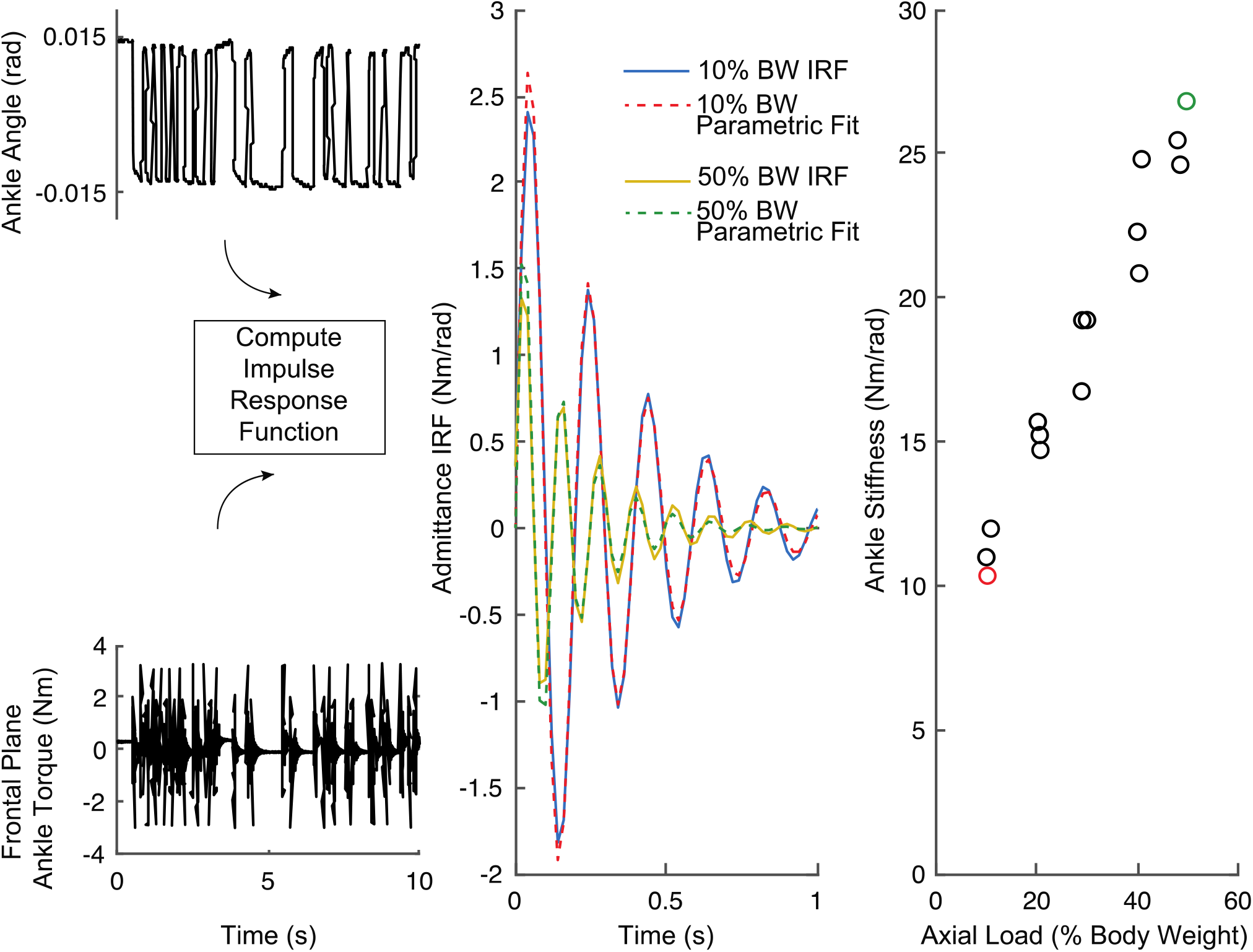
Quantifying Ankle Stiffness. The raw data (**A**), including the controlled input, ankle angle, and the measured output, frontal-plane torque, were used to compute the impulse response function defining the ankle’s impedance. (**B**) This impedance was inverted to the admittance (solid line), allowing for it to be fit to a 2^nd^ order system (dashed line). Stiffness (**C**) was computed from this parametric fit. The green and red stiffness values in (**C**) correspond to their respective green and red parametric fits in (**B**).

### EMG Analysis

For each trial, the EMG signals were notch filtered to remove any power line noise, demeaned to remove any offset, rectified, and averaged over the duration of the trial, and then subsequently normalized to the MVC of the same muscle. This analysis resulted in estimates of muscle activity as a percent of MVC for each trial, thereby allowing us to determine if there was an axial-load-dependent relationship with any observed muscle activity.

### Statistics

Our primary hypothesis was that an increase in axial load would increase ankle stiffness. We used a linear mixed-effects model to model ankle stiffness as the dependent variable, axial load as a continuous factor, sex as a fixed factor, and subject as a random factor. The same model structure was used to estimate the effect of axial load on the viscosity, inertia, and damping ratio. More details regarding the linear mixed-effects models can be found in the supplementary material. To ensure that EMG activity did not contribute to changes in stiffness over the range of applied loads, we computed a separate linear mixed-effects model for each muscle, where EMG was the dependent variable, axial load was a continuous factor, and subject was a random factor. We estimated the parameters for these models using a restricted maximum likelihood method and Satterthwaite approximations for the degrees-of-freedom (Luke, 2017). All metrics are reported as mean estimate ± standard error unless otherwise noted. For all hypothesis tests, significance levels were set to 0.05. Quality of the models were assessed by computing the R^2^, following removal of the subject-specific intercepts.

## RESULTS

Frontal-plane ankle stiffness increased linearly with an increase in axial load for all subjects. An example of one representative subject can be shown in **Figure 3**, which shows stiffness normalized by body weight (BW) increased with increasing measured axial load (as a percent of bodyweight). There was a linear increase in stiffness with the axial load from the unloaded case (about 10% of body weight) to 50% of the subject’s body weight (R^2^ = 0.98). Across all subjects (**Figure 4**), stiffness increased linearly (R^2^ = 0.96) at a rate of 0.084 ± 0.005 Nm/rad/N/%BW, (t_13_ = 15.2, p < 0.0001). As a result, at 50% BW, the ankle is approximately 3 times stiffer compared to 0% BW.

**Figure 3.**
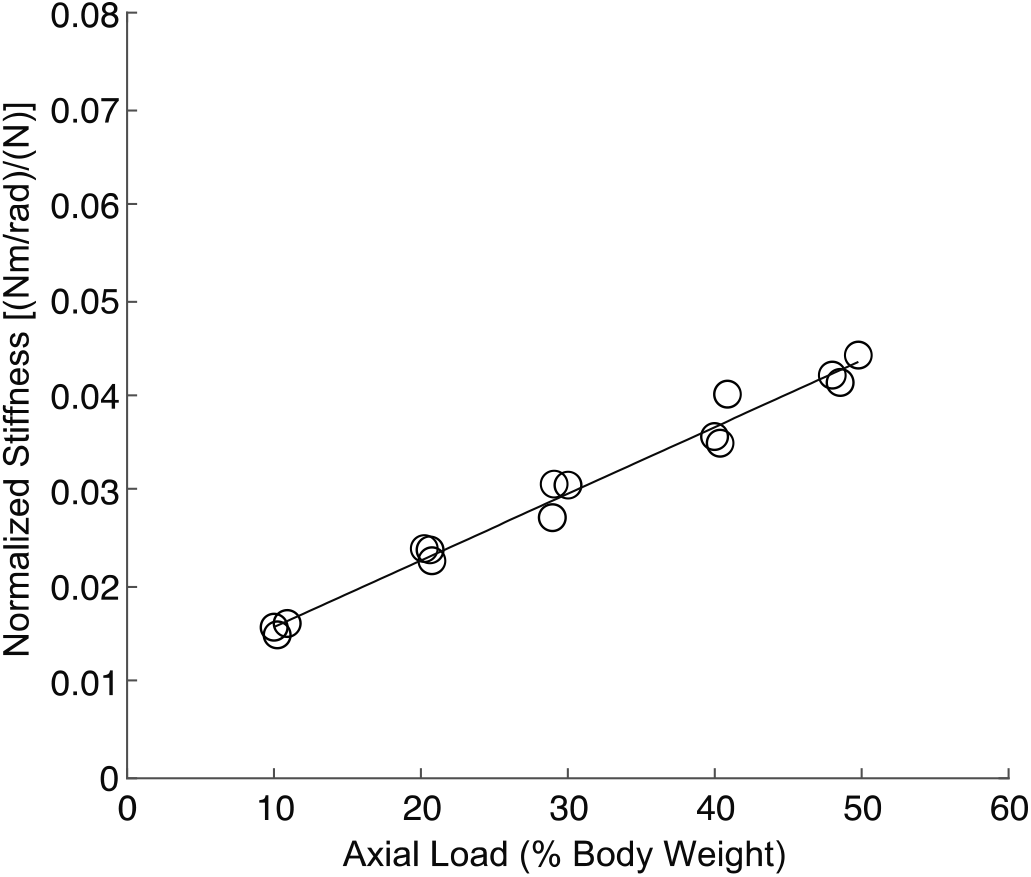
Ankle stiffness increased with axial load in a representative subject. This subject demonstrated a linear increase in stiffness across the entire range of applied load that was tested (R^2^ = 0.98). Each point shows the stiffness estimated in a trial, with the line representing a least-squares fit to the data.

**Figure 4.**
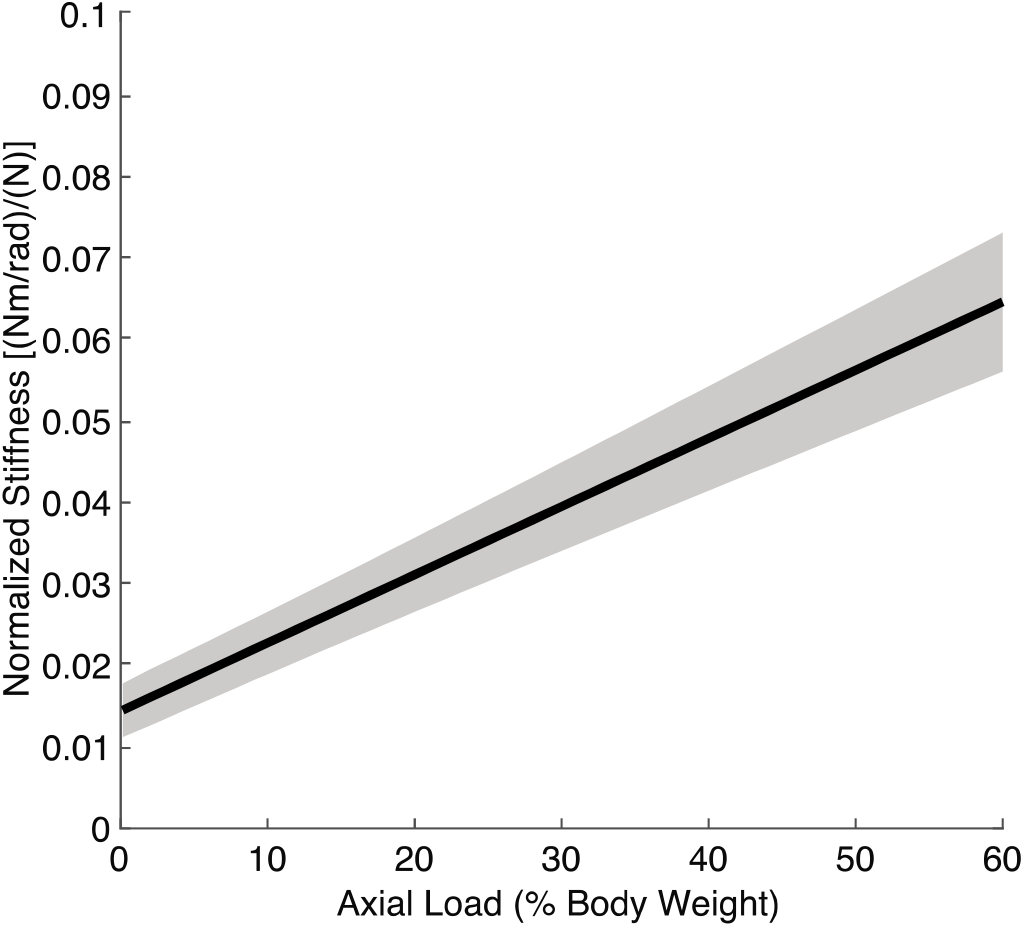
Ankle stiffness increased with axial load across all subjects. The stiffness increased linearly with axial load across all subjects (R^2^ = 0.96). The solid line shows the group average from the linear mixed-effects model, with the shaded area showing the 95% confidence intervals of the fit.

The effect of axial load on ankle stiffness was not due to an axial-load-dependent change in muscle activity. To exclude the possibility that the increase in stiffness with axial load was due to an increase in muscle activation, we monitored muscle activity throughout all the trials. **Figure 5** shows the level of activation for the 6 muscles recorded during the experiment for the same representative subject shown in **Figure 3**. For this subject, there was no muscle that surpassed an activation level of more than 5% MVC. More importantly, the level of activation did not change with the axial load applied to the ankle for any muscle. Across all subjects, there was little effect of axial load on muscle activity, with the dependence of EMG on load ranging from −0.018 to 0.034 %MVC/%BW, and none of them reaching significance (all p > 0.10, **Table 1**).

**Table 1:** Load Parameters for the Linear Mixed Effects Model for Each EMG.

| **Muscle** | **EMG - Load Slope [%MVC/%BW]** | **EMG - Load Slope Standard Error [%MVC/%BW]** | **t-statistic** | **Degrees of Freedom** | **p-value** |
| --- | --- | --- | --- | --- | --- |
| Tibialis Anterior | 0.0033 | 0.0032 | 1.03 | 35 | 0.31 |
| Lateral Gastrocnemius | 0.034 | 0.033 | 1.02 | 37 | 0.32 |
| Medial Gastrocnemius | 0.005 | 1.41 | 0.76 | 14 | 0.46 |
| Soleus | -0.018 | 0.030 | -0.61 | 56 | 0.54 |
| Peroneus Longus | 0.0073 | 0.0043 | 1.71 | 13 | 0.11 |
| Peroneus Brevis | 0.014 | 0.010 | 1.38 | 14 | 0.19 |

**Figure 5.**
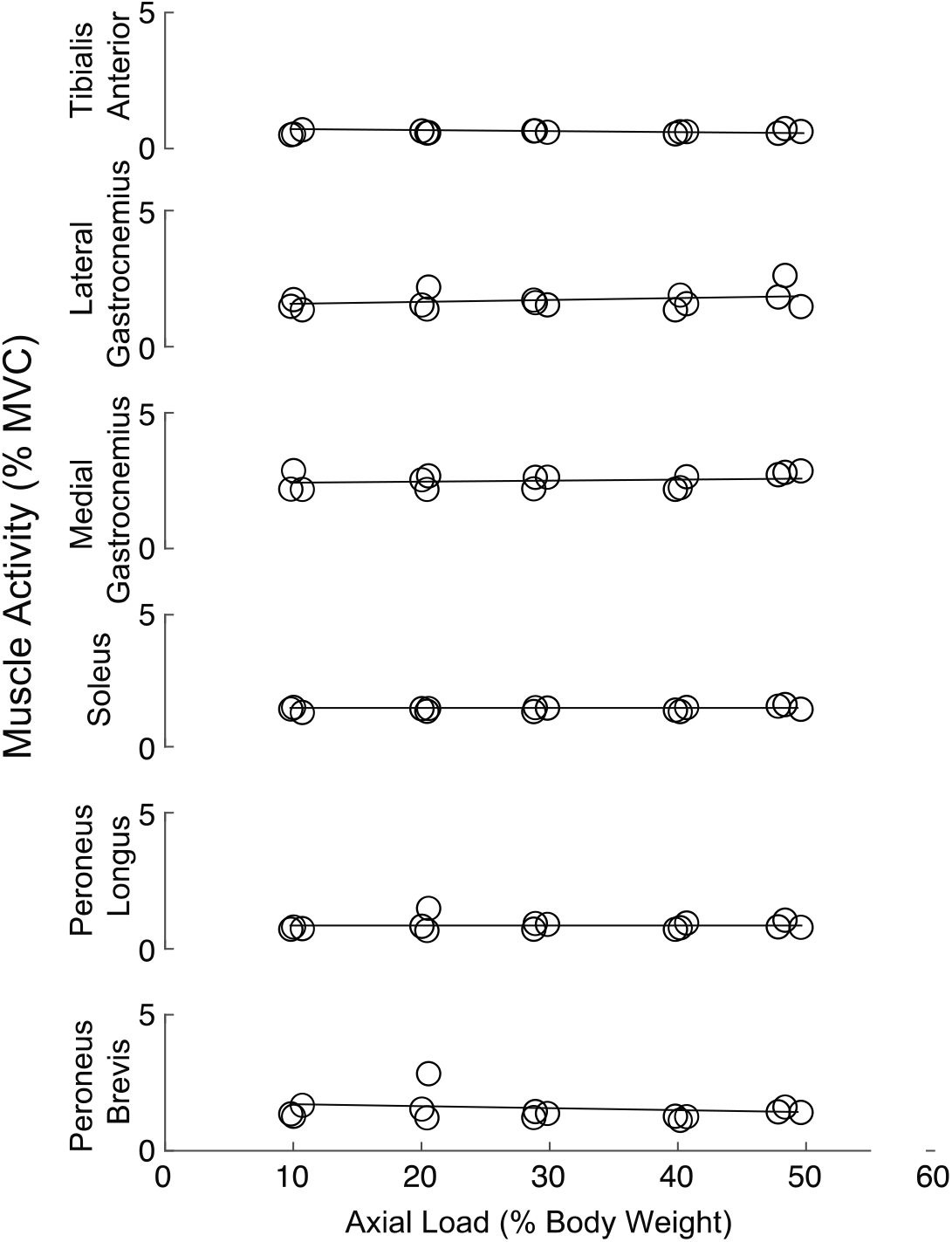
Muscle activation did not increase with load. The mean EMG is shown from the same representative subject as in **Figure 3**. The lines represent a least-squares fit to the data. All muscles remained below 5% MVC throughout the range of applied load. Muscle activation did not increase with load in any muscle. These results were consistent across the entire group, as demonstrated by the results of the linear mixed-effects model shown in **Table 1**.

Stiffness was lower in women than in men across the entire range of axial loading, but the difference did not reach significance. Stiffness was 37% reduced in females (−0.0067 +/- 0.0032 Nm/rad/N, t_13_ = −2.07, p = 0.058) at 0% axial load, while there was little difference in the rate of increase of stiffness with axial load between men and women (−0.0042 +/- 0.0106 Nm/rad/N/%BW, t_13_ = −0.40, p = 0.70, **Figure 6**).

**Figure 6.**
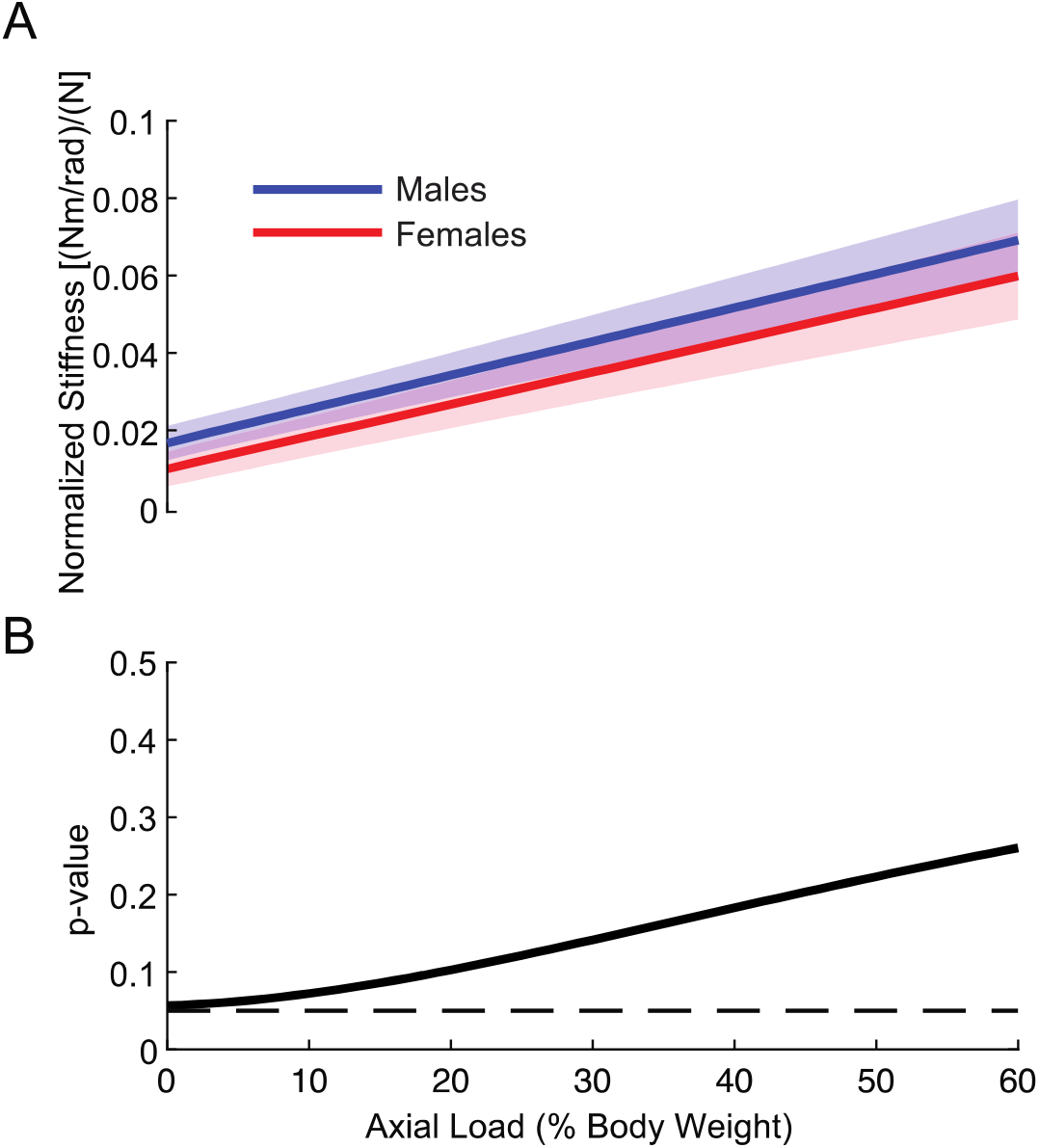
The difference in ankle stiffness between males and females did not reach statistical significance. **(A**) For both sexes, ankle stiffness increased with load applied. (**B**) Ankle stiffness was consistently higher in males than in females, but this difference did not reach significance at any of the axial load levels tested (p = 0.05, denoted by the dashed line).

While there was an increase in viscosity with axial loading, the ankle remained substantially underdamped throughout the entire range of axial loads. There was a linear increase in viscosity with axial loading (R^2^ = 0.88) at a rate of 4.5×10^−4^ ± 0.8×10^−4^ Nm/rad/s/N/%BW (t_13_ = 5.39, p = 0.0001, **Figure 7A**). As expected, inertia remained largely unchanged with axial loading 1.1×10^−6^ ± 0.7×10^−6^ kgm^2^/N/%BW (t_13_ = 1.34, p = 0.20, **Figure 7B**). Combining the estimates of stiffness, viscosity and inertia, we found that the ankle remained substantially underdamped throughout the range of axial loading, ranging from 0.05 at 0% BW to 0.21 at 50% BW (**Figure 7C**).

**Figure 7.**
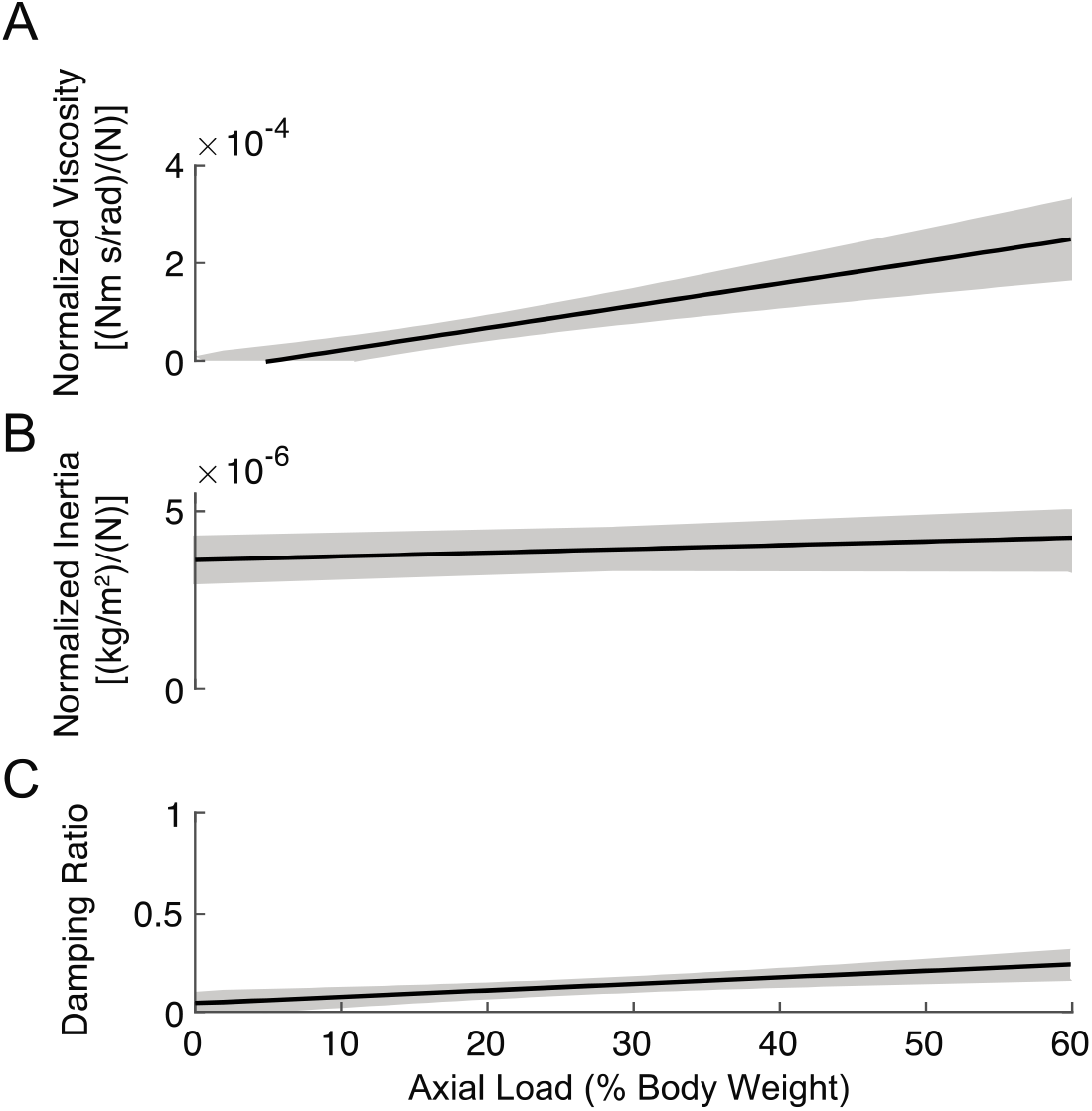
Viscosity increased with axial load, but system remained substantially underdamped at all load levels. (**A**) The viscosity increased linearly with load across the group of subjects (R^2^ = 0.88). (**B**) As expected, the inertia remained unchanged with increasing axial load (R^2^ = 0.32). (**C**) The damping remained low at all levels of load, demonstrating the minimal effect viscosity has on the ankle impedance under passive-axial-loaded conditions. The solid line in each plot shows the group average fit from the linear mixed-effects model, with the shaded area showing the 95% confidence intervals of the fit.

## DISCUSSION

The objective of this study was to determine the effect of passive axial loading on frontal-plane ankle stiffness. We accomplished this by estimating ankle stiffness in a seated configuration as axial load was applied via pressure on the knee. We tested the hypothesis that ankle stiffness would increase linearly with the applied axial load. Our hypothesis was supported as we demonstrated a significant effect of axial load on ankle stiffness. Importantly, we did not see any axial-load-dependent relationship with muscle activity, indicating that the changes in ankle stiffness were due solely to the changes in applied axial load. These results demonstrate that axial loading is a significant contributor to maintaining frontal-plane ankle stability, and disruptions to this mechanism may result in pathological cases of ankle instability.

Frontal-plane ankle stiffness increased with axial load in the seated posture. While no studies have quantified frontal-plane ankle stiffness in a seated posture with applied axial loading, a number of studies have looked at the frontal-plane ankle stiffness during weight-bearing in a standing posture. Our measures of ankle stiffness at axial loading equivalent to 50% body weight (0.055 ± 0.004 Nm/rad/N) are within the range of other studies that quantified frontal plane ankle stiffness at 50% BW [.044 – 0.112 Nm/rad/N] (A. Ribeiro et al., 2018; Matos et al., 2021; Nalam et al., 2021). Our results also agree with previous studies showing that frontal-plane ankle stiffness increased with increasing weight on the ankle during standing (Matos et al., 2021; Nalam et al., 2021). While the Matos et al. (2021) study did not attempt to isolate the effect of axial loading from the effect of muscle activation, Nalam et al. (2021) did try to isolate the effect of axial load analytically. Their technique proved reliable in the sagittal plane, but was not able to reliably predict the effect of axial loading on frontal-plane ankle stiffness. Even though these studies could not isolate the effect of axial load on frontal-plane ankle stiffness, they do provide reliable measures of how ankle stiffness increases with weight bearing (ranging from 0.0375 to 0.053 Nm/rad/N). Our estimate of the increase in stiffness with axial load is as large as those during standing, suggesting that much of the weight-bearing stiffness is due to axial loading of the ankle joint. Interestingly in our study the stiffness was actually higher than in the other studies, where muscle activation was also present. One possible explanation is that in our study the leg was constrained compared to the other studies where the entire body was unconstrained, and it is known that constraining proximal joints could lead to higher stiffness in distal joints (Perreault et al., 2000). Overall, passive axial loading appears to be an important contributor to frontal-plane ankle stiffness and likely plays a significant role in maintaining ankle stability during weight-bearing conditions.

While numerous mechanisms contribute to ankle stiffness, most of them would not be sensitive to increased passive compressive axial loading. Ligaments, muscles and tendons all contribute to the stiffness of the ankle (Leardini et al., 2000; Sakanaka et al., 2018), though all these tissues are likely under compressive loads, and the stiffness of each of these increases with tensile load not compressive load (Fung, 1993; Martin et al., 1998). Cadaveric studies suggest that the articular surfaces within the ankle joint complex may be the mechanism responsible for the increase in ankle stiffness with passive axial loading (Stiehl et al., 1993; Stormont et al., 1985; Watanabe et al., 2012). Under axial-loading conditions, the ligaments provide negligible resistance to imposed rotations in the frontal plane, implying that nearly 100% of the resistance torque to the imposed rotations is due to articular surfaces (Stormont et al., 1985). Furthermore, this resistance torque, a similar measure to the stiffness we measured, increases with axial loading (Stiehl et al., 1993; Stormont et al., 1985). This increase in resistance to imposed movements by the articular surface with increasing axial loading may be due to articular surfaces having greater articular congruity (Stiehl et al., 1993) or increased contact stress (Tochigi et al., 2006) in the load-bearing state. While our study does not allow us to directly point to a mechanism that may be contributing to the effect of axial load, our results are in line with cadaveric studies pointing to the greater contribution of articular surfaces.

While ankle stiffness was greater in men than in women, this difference did not reach significance over the range of loads tested. The observed difference arose mainly from the difference in unloaded stiffness, as there was little difference in the rate of increase of stiffness with loading between men and women. This higher stiffness in men during unloaded conditions is consistent with differences seen in the ligamentous laxity seen between men and women. During unloaded conditions, the soft tissues, such as the ligaments, are the primary contributors to ankle stability (Watanabe et al., 2012) and women have greater ligamentous laxity than men (Ericksen and Gribble, 2012; Wilkerson and Mason, 2000). While present, our results suggest that these differences are modest.

In addition to the increases in stiffness with axial loading, we also found an increase in viscosity with loading. However, this axial-loading sensitive viscosity component likely plays little role in maintaining ankle stability. Throughout the range of axial loading we tested, the ankle was substantially underdamped (no more than 0.21), demonstrating the negligible contributions of viscosity to ankle impedance. Even during the rapid dynamic conditions in which sprains occur, where we would expect viscosity to play its largest role, based on the estimates of stiffness and viscosity we computed in our study and estimates of the kinematics of ankle sprain from other studies (Fong et al., 2012), the stiffness would produce approximately ten times more resistive torque than the viscosity. Thus, axial loading increases stability of the ankle primarily by increasing the stiffness of the ankle, and not through increasing the viscosity of the ankle.

One limitation of this study is that activation of the tibialis posterior, a major invertor of the ankle, was not measured. This was due to it being a deep muscle that could not be easily recorded with surface EMG. However, we did record activation from the tibialis anterior, another inverter of the ankle and a superficial muscle. During maximum voluntary levels of isometric supination of the ankle, the tibialis anterior exhibits similar levels of activation compared to the tibialis posterior (Hagen et al., 2016). During walking, the tibialis posterior behaves similarly to the peroneus longus and the medial gastrocnemius (Murley et al., 2014), both of which we observed no load-dependent changes in activation. Based on the similarities, it is unlikely that the tibialis posterior experienced a load-dependent increase in activation while none of the other ankle muscles did.

In conclusion, we found that axial loading increases frontal-plane ankle stiffness by about 3-fold from 0% BW to 50% BW. Importantly, this is independent of muscle activation, indicating this a passive mechanism of stabilizing the ankle. These results suggest that passive axial loading of the ankle joint is an important contributor to maintaining ankle stability, and deficits in this mechanism could lead to pathological cases of ankle instability.

